# Enterovirus-A76 of South-East Asian ancestry in a Captive Chimpanzee (*Pan troglodytes*) in Jos, Nigeria

**DOI:** 10.1101/120501

**Authors:** A.O. Oragwa, U.E. George, T.O.C. Faleye, M.O. Adewumi, J.A. Adeniji

## Abstract

We recently detected EV-A119 and EV-B111 (previously shown to co-circulate between nonhuman primates (NHPs) and humans) in Nigerian children diagnosed with acute flaccid paralysis (AFP). This study was designed to investigate and catalogue EVs present in captive NHPs in Nigeria.

Twenty-seven fecal samples collected from captive NHPs in a Wild Life Park and Zoological garden at Jos, Nigeria in April 2016 were analyzed in this study. Samples were resuspended in a phosphate buffered saline (PBS)/chloroform mixture, and the clarified supernatant was subjected to RNA extraction, cDNA synthesis, a Panenterovirus 5^*I*^-UTR assay, and three different enterovirus VP1 snPCR assays. All amplicons from the snPCR assays were sequenced, and enteroviruses identified using the enterovirus genotyping tool and phylogenetic analysis.

Eight (29.63%) (two each from Chimpanzees, Patas Monkey, Mona Monkey and Baboon) of the 27 samples were positive for the 5^*I*^-UTR assay. One (3.70%) of the 27 samples was positive for the enterovirus VP1 snPCR assays in addition to its positivity by 5^*I*^-UTR assay. The same sample happens to be one of the two samples from Chimpanzees that tested positive for the 5^*I*^-UTR assay, and it was subsequently identified as EV-A76 of South-East Asia ancestry.

This study documents the first recorded attempt to detect and identify enteroviruses in NHPs in Nigeria. It also reports the first detection and identification of EV-A76 in Nigeria and particularly in a NHP. It is of utmost importance that the enterovirus VP1 assays be improved to enable detection of EVs that have been detected in NHPs but yet to be described in humans.

## Introduction

Enteroviruses are non-enveloped viruses with icosahedral symmetry, a diameter of 27–30 nm and a single-stranded, positive sense, RNA genome of ~7,500nt. They are members of the genus *Enterovirus* in the family *Picornaviridae*, order *Picornavirales*. Within the genus are twelve species which include: EV-A to -H, EV-J and *Rhinovirus A* (RV-A) to -C (http://ictvonline.org/virusTaxonomy.asp?vesion=2012). Some species are mainly populated by EVs infecting humans e.g. EV-A to -D and RV-A to -C, while others contain predominantly EVs infecting nonhuman primates (NHPs) e.g. EV-H, and EV-J. Since the establishment of a correlation between the nucleotide sequence of one of the structural genes (VP1) and serotypes (Oberste et al., 2000), it has been used for enterovirus serotype/genotype determination.

Though there has been some effort to document circulation and genetic diversity of EVs infecting African NHPs (Harvala *et al*., 2011; Sadeuh-Mba *et al*., 2014, Mombo et al., 2017) no information exist about the situation in Nigeria. Nigeria is rich in rain and mangrove forest and home to a wide range of Old World primate species, including apes (chimpanzee and gorillas) and other species of Old World monkeys. Hence, it is important to document circulation and genetic diversity of EVs infecting NHPs in Nigeria since they may harbor enteroviruses capable of cross-species transmission (Harvala *et al*., 2011; Oberste et al., 2013b; Sadeuh-Mba *et al*., 2014, Mombo et al., 2017), and even the polioviruses (Oberste et al., 2013a).

Recent findings have shown the presence of EVs which were long considered to be primarily human viruses in faecal samples collected from wild and captive NHPs (Harvala *et al*., 2011, Oberste et al., 2013a & b). This therefore suggests that NHPs may serve as reservoirs for EVs and mixing vessels for the emergence of recombinant strains. Furthermore, in Nigerian children diagnosed with acute flaccid paralysis (AFP), we recently detected EV-A119 (Adeniji et al., 2016) and EV-B111 (unpublished), which have been previously shown to co-circulates between NHPs and humans (Ayukekbong *et al*., 2013; Oberste et al., 2013a & b; Sadeuh-Mba *et al*., 2014; Harvala *et al*., 2011; Di Cristianzano et al., 2015). This therefore suggests that EVs may also co-circulate between NHPs and humans in Nigeria. This study was therefore designed to investigate and catalogue EVs present in captive NHPs in Nigeria as a first step towards describing the genetic diversity of EVs in NHPs in the country.

## Methodology

### Sample Collection and Storage

Twenty-seven fecal samples collected from NHPs in April 2016 were analyzed in this study. The samples were collected non-invasively by properly trained NHP handlers from the cages of NHPs in a Wild Life Park and Zoological garden in Jos, North-Central Nigeria. About 5g of stool was collected from freshly deposited stools on the floor of the animal cages (Table 1) and placed in a sterile container. Samples were properly labeled and transported in cold chain to the enterovirus research laboratory at the Department of Virology, College of Medicine, University College Hospital, Ibadan, Nigeria and preserved at −20°C until analysis.

**Table 1:** Profile of NHPs sampled and results of enterovirus detection and identification assays.

| S/No | Name of Animal | Scientific Name | Site of Sample Collection | No of Animals in the cage | Sex | Age (Yrs) | Age in Captivity (Yrs) | Source/ Origin | Result |  |  |
| --- | --- | --- | --- | --- | --- | --- | --- | --- | --- | --- | --- |
|  |  |  |  |  |  |  |  |  | 5'UTR | VP-1 | Serotype |
| 1 | Olive Baboon | Papio anubis | JWLP | 2 | M/F | 3 |  | Plateau |  |  |  |
| 2 | Patas Monkey | Erythrocebus patas | JZ | 1 | M | >10 | 10 |  |  |  |  |
| 3 | Patas Monkey | Erythrocebus patas | JWLP | 4 | M1/F3 | 1, 1.5, 2, 2.5 | 1, 1.5, 2, 2.5 | Bitp | Positive |  |  |
| 4 | Tantalus Monkey | Chlorocebus tantalus | JZ | 1 | F | ~10 | ~10 | Benin |  |  |  |
| 5 | Baboon | Papio anubis | JZ | 1 | M | 5 | 5 | Plateau |  |  |  |
| 6 | Chimpanzee | Pan troglodytes | JWLP | 2 | F | 8 | 8 | Bitp | Positive | Positive | EV-A76 |
| 7 | Mona Monkey | Cercopithecus mona | JWLP | 3 | M1/F2 |  |  | Plateau/ Taraba | Weak Positive |  |  |
| 8 | Chimpanzee | Pan troglodytes | JZ | 1 | F | ~54 | ~52 | Savanna/ Benin |  |  |  |
| 9 | Patas Monkey | Erythrocebus patas | JWLP | 1 | F | >3 | ~3 | Plateau |  |  |  |
| 10 | Patas Monkey | Erythrocebus patas | JZ | 2 | M | 17 | 17 | Central Africa | Weak Positive |  |  |
| 11 | Baboon | Papio anubis | JZ | 1 | F | 4 | 4 | Plateau | Positive |  |  |
| 12 | Baboon | Papio anubis | JZ | 2 | M/F | 4&6 | 4&6 |  |  |  |  |
| 13 | Chimpanzee | Pan troglodytes | JZ | 2 | M/F | 13 | 13 |  |  |  |  |
| 14 | Olive Baboon | Papio anubis | JWLP | 1 | M | 4 | 4 | Plateau | Positive |  |  |
| 15 | Tantalus Monkey | Chlorocebus tantalus | JZ | 1 | F | 20 | 20 | Benin |  |  |  |
| 16 | Mona Monkey | Cercopithecus mona | JZ | 1 | F | 10 | 6_10 | Abuja |  |  |  |
| 17 | Baboon | Papio anubis | JZ | 1 | F | 15 | 4 | Benin |  |  |  |
| 18 | Tantalus Monkey | Chlorocebus tantalus | JZ | 1 | F | >13 | 13 |  |  |  |  |
| 19 | Tantalus Monkey | Chlorocebus tantalus | JWLP | 2 | F | >1 | 1 | Plateau |  |  |  |
| 20 | Mona Monkey | Cercopithecus mona | JZ | 1 | M | 20 | 20 | Benin | Weak Positive |  |  |
| 21 | Tantalus Monkey | Chlorocebus tantalus | JZ | 2 | F | >13 | >13 |  |  |  |  |
| 22 | Chimpanzee | Pan troglodytes | JZ | 1 | M | >20 | >20 | Benin | Positive |  |  |
| 23 | Western Baboon | Papio hamadryas | JWLP | 1 | M | 7 | 7 | Lagos |  |  |  |
| 24 | Patas Monkey | Erythrocebus patas | JZ | 2 | M/F | >13 | >13 |  |  |  |  |
| 25 | Patas Monkey | Erythrocebus patas | JWLP | 1 | M | 1.5 | 1.5 | Bitp |  |  |  |
| 26 | Tantalus Monkey | Chlorocebus tantalus | JZ | 2 | M | 13_9 | 13_9 |  |  |  |  |
| 27 | Olive Baboon | Papio anubis | JWLP | 1 | M | 7 | 7 | Plateau |  |  |  |

NB: JWLP= Jos Wild Life Park., JZ= Jos Zoo, Bitp= Born in the park

### Sample processing

About one gram of each stool specimen was diluted in 3mL phosphate buffered saline (PBS), one milliliter chloroform and one gram of glass beads. The mixture was vortexed for two (2) minutes and subsequently centrifuged at 3000 rpm for 20 minutes. Afterwards, 2mL of the supernatant was aliquoted in one milliliter volumes into two cryovials per sample. One vial was stored at −20°C while the other was analyzed further.

### RNA Extraction and cDNA Synthesis

RNA was extracted from 100μL of each of the 27 samples using Jena Bioscience RNA extraction kit (Jena Bioscience, Jena, Germany) and was used for cDNA synthesis using Script cDNA Synthesis Kit (Jena Bioscience, Jena, Germany) as previously described (Adeniji and Faleye, 2014; Faleye et al., 2016a). To be precise, two different cDNA (one with random hexamers [Adeniji and Faleye, 2014], and the other with primers AN32-AN35 [Faleye et al., 2016a]) were made in 10μL volumes, for all the samples. This mixture was incubated at 42 °C for 10min followed by 50 °C for 60 minutes in a Veriti thermocycler (Applied Biosystems, California, USA).

## Polymerase chain reaction (PCR) Screens

### Polymerase chain reaction (PCR) assay for the 5^*I*^ UTR

The 5^*I*^-UTR assay was done in 30μL volume. The cDNA made with random hexamers was used for this assay. Every PCR tube contained 10μL of cDNA, 6μL of Red Load Taq (Jena Bioscience, Jena, Germany), 13.4μL of RNase free water and 0.3μL each of the forward (PanEnt-5^*I*^-UTR-F) and reverse (PanEnt-5^*I*^-UTR-R) primers (Adeniji and Faleye, 2014). A Veriti thermal cycler (Applied Biosystems, California, USA) was used for thermal cycling as follows; 94^0^C for 3 minutes, then 45 cycles of 94°C for 30 seconds, 42°C for 30 seconds, and 60°C for 30 seconds, with ramp of 40% from 42°C to 60°C. This was then followed by 72°C for 7 minutes, and held at 4°C until the reaction was terminated. Subsequently, PCR products were resolved on 2% agarose gel stained with ethidium bromide and viewed using a UV transilluminator.

### Seminested Polymerase chain reaction (snPCR) assay for the VP1 gene

As previously described (Faleye et al., 2016a & b), an snPCR assay was done in this case. The assay is a modification (Faleye et al., 2016a & b) of the WHO recommended RT-snPCR assay for direct detection of enteroviruses from clinical specimen (WHO, 2015). A single first round PCR was done. Followed by three (PanEnterovirus [PE], Enterovirus Species A or C [EV-A/C] and Enterovirus Species B [EV-B]) different second round PCR assays. The cDNA made with primers AN32-AN35 was used for this assay. Each reaction of the first round assay was also done in 30μL volume and contained 10μL of cDNA, 6μL of Red Load Taq (Jena Bioscience, Jena, Germany), 13.4μL of RNase free water and 0.3μL each of the forward (224) and reverse (222) primers. For the three second round PCR assays, the reaction mix was constituted as above except for the cDNA that was substituted with the first-round PCR product. Also, instead of using 10μL of cDNA, 3μL of first round PCR product was used, and the RNase free water was increased to 20.4μL. In addition, the primers were also changed. Primer AN88 was reverse primer used for all three second round assays. However, the forward primers were AN89, 189 and 187 for PE, EV-A/C, EV-B assays, respectively.

The same Veriti thermal cycler (Applied Biosystems, California, USA) used for the 5^*I*^-UTR assay was also used for the VP1 snPCR assay. Thermal cycling conditions were as described for the 5^*I*^-UTR assay but with few changes. The extension times were 60 and 30 seconds for the first and second round assays respectively. All PCR products were also resolved on 2% agarose gel stained with ethidium bromide and viewed using a UV transilluminator.

## Amplicon Sequencing and Enterovirus Identification

Only amplicons with expected band size emanating from the second round PCR assays for the VP1 gene were shipped to Macrogen Inc, Seoul, South Korea, for purification and nucleotide sequencing using the respective primer pairs. The enterovirus genotyping tool (EGT) (Kroneman et al., 2011) was subsequently used for enterovirus type determination.

## Phylogenetic analysis

For multiple sequence alignment of the sequence(s) generated in this study and those of reference sequences downloaded from Genbank, the CLUSTAL W program in MEGA 5 software (Tamura *et al*., 2011) was used with default settings. A neighbor-joining tree was then constructed using the Kimura-2 parameter model (Kimura, 1980) and 1,000 bootstrap replicates in the same MEGA 5 software.

**Nucleotide Sequence Accession Numbers:** The sequence(s) generated in this study have been deposited in GenBank under accession numbers KY798125

## RESULTS

### Polymerase chain reaction (PCR) assay for the 5^*I*^ UTR

Eight (29.63%) of the 27 samples subjected to this assay had the expected band size (~114bp). The eight bands were detected in samples from four different NHP types namely, Chimpanzees, Patas Monkey, Mona Monkey and Baboon (Table 1). It is however crucial to note that while five (5) of the bands had very high intensity, the remaining three (3) bands were very weak.

### Seminested Polymerase chain reaction (snPCR) assay for the VP1 gene

Only one of the 27 (3.70%) samples subjected to these three assays had the expected band size (~350bp). The sample was positive for all three second round PCR assays (Table 1). It is one of the two samples from Chimpanzees that had been previously positive for the 5^*I*^-UTR PCR assay.

### Amplicon Sequencing and Enterovirus Identification

The EGT independently identified all three sequences derived from the three amplicons generated from VP1 assay positive sample as Enterovirus A76 (EV-A76).

### Phylogenetic analysis

Phylogenetic analysis showed that the EV-A76 detected in this study is not related to all EV-A76 strains previously detected in sub-Saharan Africa from 2003 till date. Rather it is a member of one of the EV-A76 lineages circulating in South-East Asia and shares a common ancestor with a strain recovered in 2009 from the stool of a child in India.

## DISCUSSION

To the best of the authors’ knowledge, this study documents the first recorded attempt to detect and identify enteroviruses in NHPs in Nigeria. Here we report the first detection and identification of EV-A76 in Nigeria and particularly in a NHP. Studies (Kumar et al., 2012; Rao et al., 2012; Patil et al., 2015) have documented the recovery of EV-A76 in humans with different clinical conditions ranging from gastroenteritis (Patil et al., 2015), to central nervous system (CNS) manifestations like acute flaccid paralysis (AFP) (Rao et al., 2012; Sadeuh-Mba et al., 2013) and encephalitis (Kumar et al., 2012). Hence, EV-A76s potential to have serious clinical manifestations in humans is obvious. Also crucial is the fact that EV-A76 has also been previously recovered in NHPs in Cameroon (Harvala *et al*., 2011; Sadeuh-Mba et al., 2014), and Bangladesh (Oberste et al., 2013a). In fact, it was unambiguously shown (Oberste et al., 2013a) and confirmed in this study (Figure 1) that in 2008, the same strain of EV-A76 was simultaneously circulating in both humans and NHPs in Bangladesh. This confirms the zoonotic potential of EV-A76 and lays more credence to the need to better understand its epidemiology.

**Figure 1:**
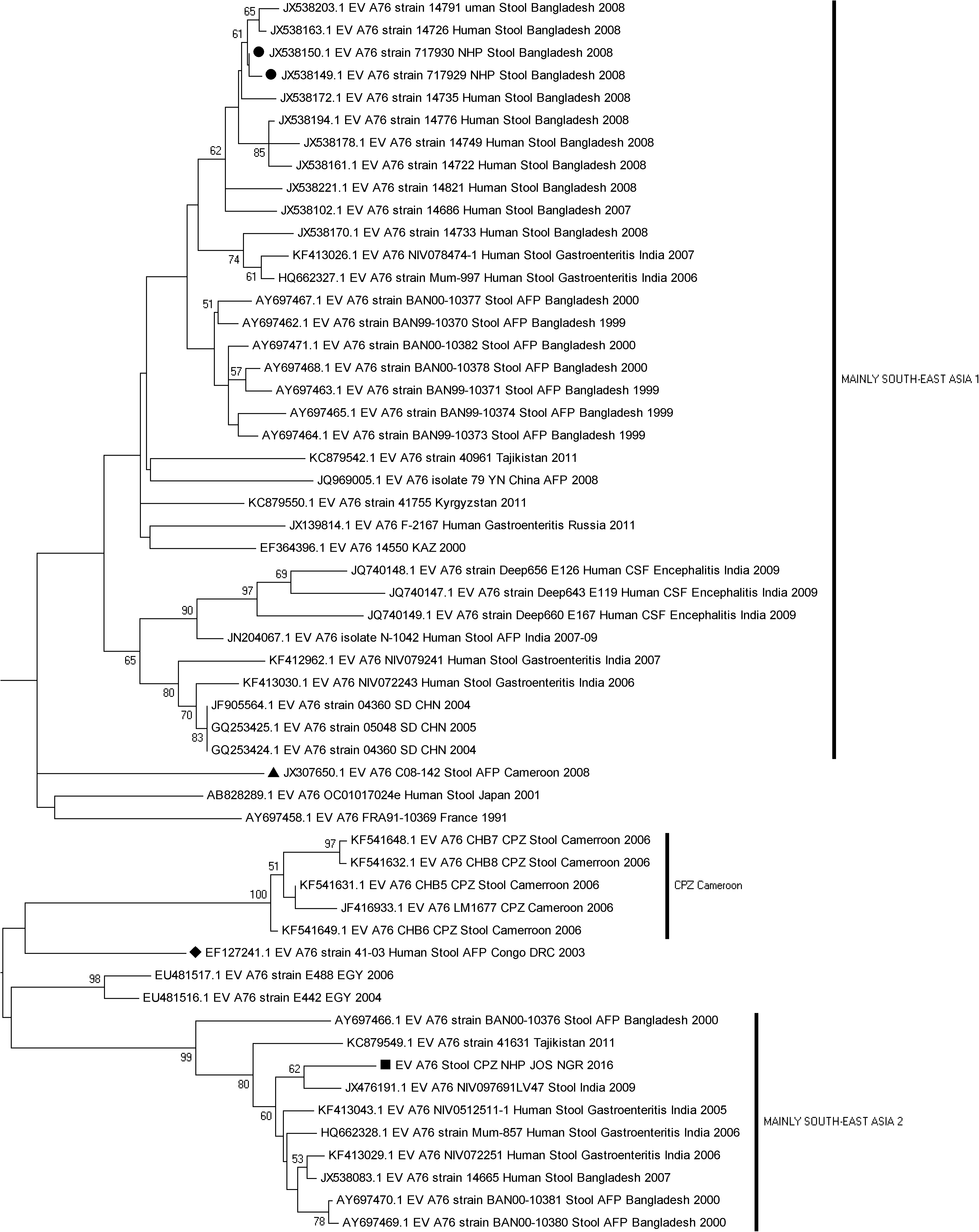
Phylogram of EV-A76 identified in this study. The phylogram is based on an alignment of partial VP1 sequences. The newly sequenced strain is highlighted with Black Square. Strains recovered from children with AFP in Congo DRC and Cameroon are indicated with black diamond and triangle, respectively. Strains recovered from NHPs in Bangladesh are highlighted with black circle. All strains recovered from NHPs in Cameroon form a cluster and are highlighted as ‘CPZ CAMEROON’ in the Phylogram The GenBank accession number of the strains are indicated in the phylogram. Bootstrap values are indicated if > 50%.

Prior this finding of EV-A76 in a Chimpanzee in Nigeria, it had been previously described in two other countries in sub-Saharan Africa; Cameroon (Harvala *et al*., 2011; Sadeuh-Mba et al., 2013; 2014) and Congo DRC (Junttila et al., 2007). It is however important to emphasize that the EV-A76 strain detected in this study is not related to those previously detected in the sub-region. Rather, it belonged to one of the lineages that had been circulating in South-East Asia since year 2000. In fact, the EV-A76 strain detected in this study is most closely related to (shares a common ancestor) a strain detected in India in 2009. How the infected Chimpanzee got exposed and infected by this imported strain of EV-A76 is currently not clear. However, considering the Chimpanzee in question was born in the park (Table 1), and had been in captivity in the same park since birth, it must have been exposed to this EV-A76 strain at the park. The possibility therefore exists that this strain of EV-A76 might be circulating either in the park or the encompassing community. In fact, the topology of the relationship between the India 2009 strain and that detected in this study suggests that several isolates of EV-A76 that are similar to both are yet to be described. Hence, there is need for further investigation to better describe and catalogue the diversity of EV-A76 in the region and Nigeria at large.

In this study, the 5^*I*^-UTR PCR assay detected enteroviruses in 29.63% (8/27) of the samples analyzed. However, the VP1 gene was amplified in only one of the eight samples positive for the 5^*I*^-UTR PCR assay. The gene was unamplifyable in the remaining seven samples. Why this is the case is currently not clear. However, as previously suggested (Mombo et al., 2017), the possibility exists that the VP1 gene could not be amplified in the other seven samples because they are not members of EV-A – EV-D for which the VP1 amplification snPCR assay was originally developed (Nix et al., 2006). With the increased significance of enteroviruses in NHPs, it is of utmost importance that the WHO recommended assay (WHO, 2015) be further improved to be able to consistently amplify the VP1 gene of non-Species A – D enteroviruses.

## ACKNOWLEDGEMENTS

We thank the management and staff of both the Wild Life Park and the Zoo in Jos, Nigeria where the samples analyzed in this study were collected. We also thank the onsite animal handlers who helped with collection of all the samples.

## AUTHOR CONTRIBUTIONS

1. Study Design (All authors)
2. Sample Collection, laboratory and Data analysis (All authors)
3. Wrote, revised, read and approved the final draft of the Manuscript (All authors)

